# An Approach for Estimating Explanation Uncertainty in fMRI dFNC Classification

**DOI:** 10.1101/2022.05.23.493148

**Authors:** Charles A. Ellis, Robyn L. Miller, Vince D. Calhoun

**Affiliations:** Tri-institutional Center for Translational, Research in Neuroimaging and Data Science, Georgia State University, Georgia Institute of Technology, and Emory University, Atlanta, USA

**Author notes:** Funding is provided by NIH R01MH118695 and NSF 2112455.

**Keywords:** Explainable AI, explanation uncertainty, fMRI, functional network connectivity, convolutional neural networks

## Abstract

In recent years, many neuroimaging studies have begun to integrate gradient-based explainability methods to provide insight into key features. However, existing explainability approaches typically generate a point estimate of importance and do not provide insight into the degree of uncertainty associated with explanations. In this study, we present a novel approach for estimating explanation uncertainty for convolutional neural networks (CNN) trained on neuroimaging data. We train a CNN for classification of individuals with schizophrenia (SZs) and controls (HCs) using resting state functional magnetic resonance imaging (rs-fMRI) dynamic functional network connectivity (dFNC) data. We apply Monte Carlo batch normalization (MCBN) and generate an explanation following each iteration using layer-wise relevance propagation (LRP). We then examine whether the resulting distribution of explanations differs between SZs and HCs and examine the relationship between MCBN-based LRP explanations and regular LRP explanations. We find a number of significant differences in LRP relevance for SZs and HCs and find that traditional LRP values frequently diverge from the MCBN relevance distribution. This study provides a novel approach for obtaining insight into the level of uncertainty associated with gradient-based explanations in neuroimaging and represents a significant step towards increasing reliability of explainable deep learning methods within a clinical setting.

## 1. Introduction

Explainability methods have been used extensively within neuroimaging classification [1]. These methods give insight into the key features learned by classifiers and have implications for the future use of neuroimaging classifiers in healthcare. Unfortunately, it is difficult to estimate the reliability of any explanations that may be obtained or to know the degree to which explanations can be trusted [2]. In this study, we present a novel approach that integrates existing explainability methods [3] with a method for obtaining Bayesian-like uncertainty estimates for model predictions [4] to obtain a better understanding of the reliability of neuroimaging explanations.

Some methods have been developed that could be used for estimating the degree of confidence in an explanation. These methods chiefly involve repeatedly perturbing data samples and examining the effect of the perturbations on model performance or classification probabilities [5], [6]. Unfortunately, the utility of perturbation approaches can be reduced by high dimensional data spaces [7], and perturbation approaches can produce out-of-distribution samples that make explanations unreliable [8]. An alternative approach would involve repeatedly retraining a model on the same data and outputting explanations for each new model. This approach could be used with gradient methods that would be able to provide explanations for highly dimensional data. However, this would be very computationally expensive and have little practical utility in an application setting. Lastly, some methods have been developed with the goal of determining whether explanations are valid [9]. However, they do not really indicate a level of confidence in an explanation, and they can be computationally intensive.

In recent years, multiple methods for obtaining Bayesian-like uncertainty estimates for predictions from non-Bayesian neural networks have been developed. One of these methods, Monte Carlo dropout [10], has been used extensively in the domain of neuroimaging classification and has demonstrated the ability to improve model test performance in some instances [11]. Monte Carlo dropout involves using dropout during model training and testing such that the model can be tested many times with different initializations of dropout. At each initialization, a separate prediction can be generated, giving an approximately Bayesian distribution of predictions. Unfortunately, the use of dropout during testing can, in some cases, destroy model gradients, which makes Monte Carlo dropout poorly compatible with gradient-based explainability methods. An alternative approach is Monte Carlo batch normalization (MCBN) [4], which does not adversely affect model gradients and is relatively novel to the domain of neuroimaging classification.

In this study, we seek to combine explainability methods with methods for estimating prediction uncertainty to obtain insight into the degree of certainty associated with explanations. We train a 1-dimensional convolutional neural network (1D-CNN) to discriminate between individuals with schizophrenia (SZ) and healthy controls (HC) using dynamic functional network connectivity (dFNC) time-series extracted from resting-state functional magnetic resonance imaging (rs-fMRI). We implement MCBN for what is, to the best of our knowledge, its first use with neuroimaging data to improve model generalization. We then integrate the MCBN with layer-wise relevance propagation (LRP) [3] to obtain insight into the uncertainty associated with the explanations.

## II. Methods

### A. Description of dataset

We used rs-fMRI data from the Functional Imaging Biomedical Informatics Research Network (FBIRN). It has been used in many fMRI studies [7], [12]. It includes 151 SZs and 160 HCs collected from the University of California at Irvine, the University of California at Los Angeles, the University of California at San Francisco, Duke University, the University of North Carolina at Chapel Hill, the University of New Mexico, the University of Iowa, and the University of Minnesota. Informed consent and Institutional Review Board approval was obtained at each site. One 3T General Electric and six 3T Siemens scanners were used for collection. An AC-PC aligned echo-planar imaging sequence with TR=2s, TE=30ms, flip angle=77°, voxel size= 3.4×3.4×4mm^3^, slice gap=1mm, 162 frames, and 5:24 minutes was used to collect T2*-weighted functional images. Eyes were closed during recording.

### B. Description of Preprocessing

Statistical parametric mapping (https://www.fil.ion.ucl.ac.uk/spm/) was used for preprocessing. We accounted for head movement via rigid body motion correction. We performed spatial normalization to an echo-planar imaging template in standard Montreal Neurological Institute space and resampled to 3×3×3 mm^3^.We then smoothed the images via a Gaussian kernel (full width at half maximum = 6mm). We then used the Neuromark automatic independent component analysis pipeline in the GIFT toolbox to extract 53 independent components (ICs). The 53 ICs were separated into 7 networks: subcortical (SCN), auditory (ADN), sensorimotor (SMN), visual (VSN), cognitive control (CCN), default mode (DMN), and cerebellar (CBN). To calculate dFNC values, we used a sliding window approach with a tapered window formed by convolving a rectangle (window size = 40s) with a Gaussian (σ=3). Pearson’s correlation was calculated between each of the 53 ICs at each time step. This yielded a matrix of 1378 dFNC features x 124 time points for each subject. The 1378 dFNC features can be divided into 28 network domain pairs. An example abbreviation of the connectivity between SCN and ADN is SCN/ADN.

### C. Model Development

We developed a 1D-CNN architecture shown (Figure 1). Before training, we z-scored each dFNC feature across all samples. We then used a 10-fold stratified shuffle split cross-validation approach with around 80%, 10%, and 10% of subjects assigned to training, validation, and test folds. To account for the number of features and small number of samples, we tripled the training data size in each fold by adding randomly generated Gaussian noise (μ = 0, σ = 0.7) to two copies of each training sample. We used an Adam optimizer with an initial learning rate of 0.001 that decreased by 50% after each 15 epochs that did not have a corresponding increase in validation accuracy. We used class-weighted categorial cross-entropy to account for class imbalances, and we used 100 epochs with shuffling and a batch size of 50. When testing, we used model checkpoints to select the model from the epoch that had the highest validation accuracy. We then calculated the mean (μ) and standard deviation (σ) of the sensitivity (SENS), specificity (SPEC), and balanced accuracy (BACC) metrics.

**Figure 1.**
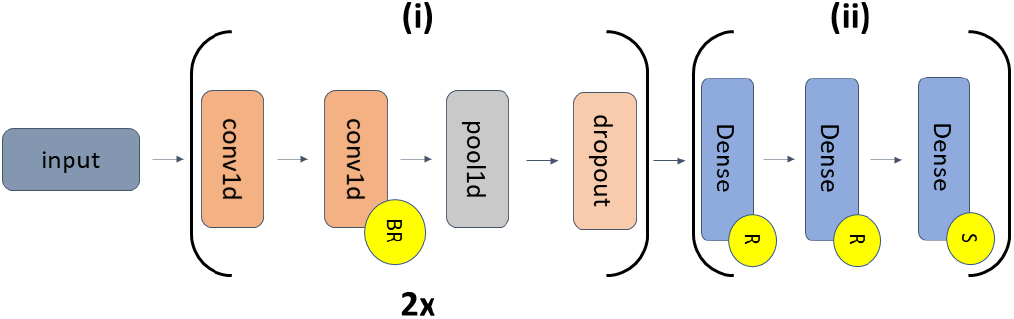
CNN Architecture. The model has feature extraction (i) and classification (ii) layers. (i) repeats twice with different hyperparameters. The first and second pairs of convolutional (conv1d) layers (kernel size = 10) have 16 and 24 filters, respectively. They are followed by a max pooling layer (pool size = 2) and spatial dropout (rates = 0.3 and 0.4). (ii) has 3 dense layers with 10, 6, and 2 nodes. Kaiming He normal initialization was used. Yellow circles with “BR”, “R”, and “S” indicate layers with batch normalization and ReLU, ReLU, and softmax activations, respectively.

### D. Monte Carlo Batch Normalization

MCBN was first presented in [4]. It draws on the idea that when batch normalization is used different minibatches of training data and the updated batch normalization layer parameters that results from model training with each minibatch represents a Bayesian process where the different randomly generated minibatches represent a prior. In our implementation of MCBN, we trained the model. We then iteratively generated minibatches of training data and updated the batch normalization parameters with the loss from each minibatch. After updating parameters at each iteration, we generated predictions and explanations for the test data. We then repeated the process for 100 minibatches.

### E. Explainability Analysis

We used LRP to generate explanations for the CNN without MCBN and during each iteration of MCBN. We used the CNN from the fold with the best BACC. Originally developed for image classification, LRP has since been used in multiple neuroscience studies involving time-series [1], [13], [14]. LRP assigns a total relevance of 1 to the output node associated with a sample. It then propagates portions of that total relevance through each layer of the network using relevance rules such that the total relevance at each layer sums to 1. In this study, we used the αβ-rule (α=1, β=0), which only propagated positive relevance (evidence for a sample belonging to the class to which it was assigned by the model). After propagating the relevance through the network to the sample space, we summed the total absolute relevance of each dFNC node across all time steps. We also normalized the relevance for each subject such that its total absolute relevance summed to 100.

We performed several statistical analyses comparing differences in relevance between SZs and HCs and between values with and without MCBN. (1) We summed the total normalized absolute relevance in each network pair for each MCBN batch for each subject. We then used a Kolmogorov-Smirnov test to determine whether the relevance in each network domain pair was normally distributed. For those that were and were not normally distributed, we performed two-tailed t-tests and rank sum tests, respectively, to determine whether there were significant differences in MCBN relevance between SZs and HCs. We then applied FDR correction (p < 0.05) to reduce the likelihood of false positives. We repeated this for the mean MCBN relevance across iterations and for relevance values from all MCBN iterations and without MCBN. (2) To test whether each regular LRP value represented the MCBN distribution for the same node, we used Wilcoxon ranked sum tests. We did this for each subject and then calculated the percent of subjects that had significant changes (p < 0.05) in relevance of each node.

## III. Results and discussion

Here, we describe and discuss our model performance and explainability results.

### A. Model Performance

Table 1 shows μ and σ of the model performance with and without MCBN. Performance was well above chance-level, having mean values greater than 75%. Performance without MCBN favored SENS over SPEC. MCBN had beneficial effects on performance. It largely balanced the disparity between the SENS and SPEC metrics while also slightly increasing BACC.

**Table 1.** Classifier Performance

|  | SPEC | SENS | BACC |
| --- | --- | --- | --- |
| **Original** | 73.12±12.52 | 77.50±10.90 | 75.31±07.58 |
| **MCBN** | 76.25±11.79 | 75.63±12.33 | 75.94±07.79 |

### B. Explainability Results with MCBN

Figure 2 shows the mean of the μ and σ of the percent of overall relevance assigned to each node for SZs and HCs. Based on visual inspection, there were multiple noteworthy differences in the relevance distributions of SZs and HCs. For intra-network connectivity, SZs had more relevance in the SCN/SCN, VSN/VSN, and CBN/CBN, which also coincided with increases in σ. Multiple differences also occurred in inter-network connectivity relevance. Specifically, the interaction of the thalamus in the SCN with the CCN was more important for HCs than SZs. The interaction of the postcentral gyrus of the SMN with the VSN was more important in SZs than HCs. Additionally, the interaction of the hippocampus in the CCN with the CBN was more important for HCs than SZs.

**Figure 2.**
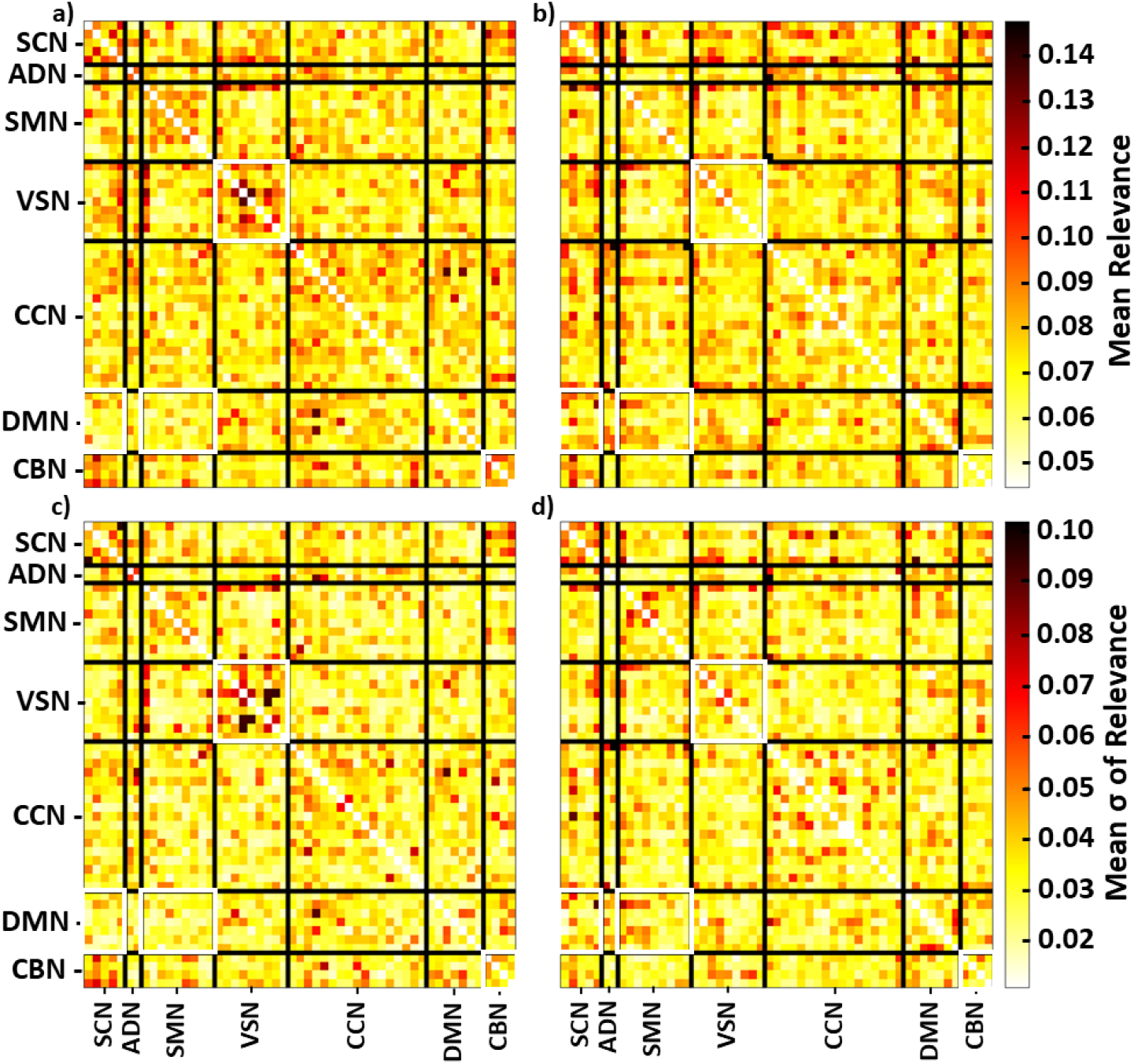
LRP Relevance for Each Class. Panels a) and b) show the average of the mean relevance for SZs and HCs across MCBN iterations, respectively. Panels c) and d) show the average of the relevance standard deviations for SZs and HCs across MCBN iterations, respectively. Each FNC domain is separated by black lines, and the color bars to the right of panels b) and d) indicate the values associated with the heatmap. The x- and y-axes show each rs-fMRI network. White boxes around individual FNC domains indicate those that had significant uncorrected differences between HCs and SZs in their mean total relevance.

Figure 2 shows the μ of relevance and thus does not account for the variation in relevance across MCBN iterations. When we tested for differences in the μ total relevance in each network domain pair across MCBN iterations, we found only four pairs – VSN/VSN, DMN/SCN, DMN/SMN, and CBN/CBN – with significant differences without FDR correction and no network pairs with significant differences after correction. In contrast, when we tested for differences between the total relevance in each network domain pair between SZs (100 iterations × 16 test participants = 1600 samples) and HCs (100 iterations × 16 test participants = 1600 samples), we found significant differences in all pairs except CCN/ADN and DMN/VSN (p < 0.05).

### C. Difference in Explanations With and Without MCBN

While MCBN increased the likelihood of identifying inter-class differences in relevance, we also wanted to determine how much MCBN affected subject-level relevance. Figure 3 shows the result of comparing whether the regular LRP relevance for each subject was away from the median MCBN-based relevance of each node. We found that MCBN affected 12% - 18% of SZs and HCs across many dFNC nodes. Interestingly, more HCs seemed to have off-median regular LRP values This was especially the case for many CCN network pairs (i.e., CCN/CCN, VSN/CCN, and SCN/CCN). These results indicate that MCBN-based explanations affect the relevance assigned to nodes on a per-subject basis and that regular LRP provides off-median relevance values for HCs more often than SZs, which has implications for the clinical use of explainability methods.

**Figure 3.**
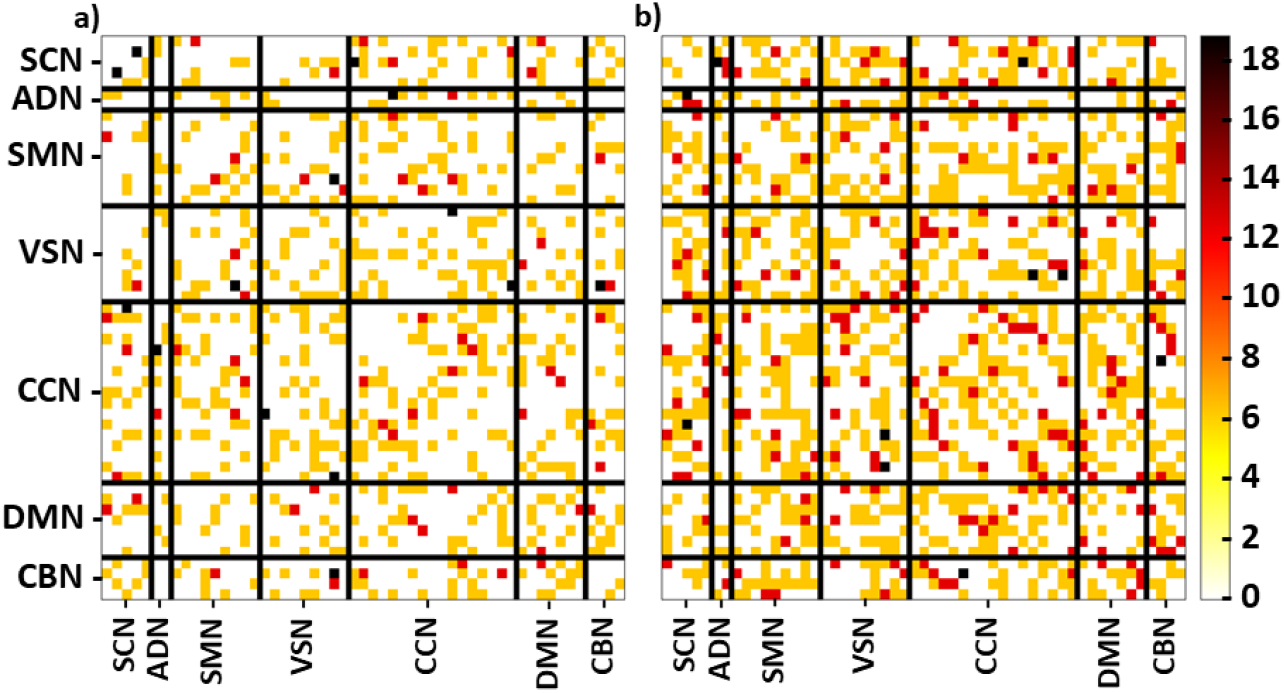
Percent of Samples with Change in Relevance Distribution. Panels a) and b) show, for each node-pair, the percent of SZ and HC subjects respectively in which traditional LRP relevance differed significantly from the center of the MCBN-based relevance distributions. Each functional domain is separated by black lines, and the color bar to the right indicates the percentages associated with the heatmap. The x- and y-axes are associated with each rs-fMRI network.

### D. Limitations and Next Steps

Our analysis focused on the effects of MCBN on the levels of importance for individual nodes and network connectivity pairs and thus involved the summation of relevance for each node across all time points. This summation could have obscured important aspects of the MCBN LRP values, and future analyses will be needed to investigate this possibility. Additionally, it might be interesting to explore the effects of MCBN upon other gradient-based explainability methods and upon explanations for other forms of neuroimaging data.

## IV. Conclusion

In this study, we introduced a novel approach for estimating the uncertainty associated with explanations by combining MCBN with LRP. We applied our approach to a CNN trained on rs-fMRI dFNC data from SZs and HCs. We identified differences in the explanations for SZs and HCs, and we found that traditional LRP explanations were in many instances away from the center of the distribution of MCBN LRP explanations. We will further develop this approach in the future in the context of other explainability methods and expect it to lead to a better understanding of the reliability of explainability methods.

## Acknowledgment

We thank those who collected the FBIRN dataset.

## Notes

### Competing Interest Statement

The authors have declared no competing interest.

### Summary of Updates

We found a minor error in the analysis and have corrected that in the updated version. It does not fundamentally affect the findings or contributions of the paper.

